# Emergence of division of labor in tissues through cell interactions and spatial cues

**DOI:** 10.1101/2022.11.16.516540

**Authors:** Miri Adler, Noa Moriel, Aleksandrina Goeva, Inbal Avraham-Davidi, Simon Mages, Taylor S Adams, Naftali Kaminski, Evan Z Macosko, Aviv Regev, Ruslan Medzhitov, Mor Nitzan

**Affiliations:** Broad Institute of Massachusetts Institute of Technology and Harvard, Cambridge, MA, USA; Tananbaum Center for Theoretical and Analytical Human Biology, Yale University School of Medicine, New Haven, CT, USA; School of Computer Science and Engineering, The Hebrew University of Jerusalem, Jerusalem, Israel; Gene Center and Department of Biochemistry, Ludwig-Maximilians-University Munich, Munich, Germany; Section of Pulmonary, Critical Care and Sleep Medicine, Yale University School of Medicine, New Haven, CT, USA; Massachusetts General Hospital, Department of Psychiatry, Boston, MA, USA; Department of Biology, Massachusetts Institute of Technology, Cambridge, Massachusetts, USA; Genentech, 1 DNA Way, South San Francisco CA, USA; Howard Hughes Medical Institute, Department of Immunobiology, Yale University School of Medicine, New Haven, CT, USA; Racah Institute of Physics, The Hebrew University of Jerusalem, Jerusalem, Israel; Faculty of Medicine, The Hebrew University of Jerusalem, Jerusalem, Israel

## Abstract

Most cell types in multicellular organisms can perform multiple functions. However, not all functions can be optimally performed simultaneously by the same cells. Functions incompatible at the level of individual cells can be performed at the cell population level, where cells divide labor and specialize in different functions. Division of labor can arise due to instruction by tissue environment or through self-organization. Here, we develop a computational framework to investigate the contribution of these mechanisms to division of labor within a cell-type population. By optimizing collective cellular task performance under trade-offs, we find that distinguishable expression patterns can emerge from cell-cell interactions *vs*. instructive signals. We propose a method to construct ligand-receptor networks between specialist cells and use it to infer division-of-labor mechanisms from single-cell RNA-seq and spatial transcriptomics data of stromal, epithelial, and immune cells. Our framework can be used to characterize the complexity of cell interactions within tissues.

## Introduction

Many cell types in multicellular organisms are multi-functional: for example, epithelial cells perform sensory, secretory, transport and defense functions. Moreover, some functions, such as transport and defense functions in epithelial cells, are bound by trade-offs: they cannot be optimized at the same time in the same cells. Division of labor is a common strategy to handle such functional trade-offs, but how it occurs within a given cell type population remains largely unknown. Cells’ expression profiles provide an opportunity to characterize division of labor in tissues, to infer the functional constraints driving it, and to determine its underlying mechanisms.

In particular, low-dimensional representations of patterns of cellular gene expression (Figure 1A) (“gene expression space”), can reflect not only distinct cell types as discrete clusters and dynamical processes, such as cell differentiation along continuous trajectories (Wagner, Regev, and Yosef 2016; Tanay and Regev 2017; Sagar and Grün 2020; Ding, Sharon, and Bar-Joseph 2022), but also the collective optimization of task performance under trade-offs within a given cell type (Shoval et al. 2012; Korem et al. 2015; Hart et al. 2015; Hausser et al. 2019). A theoretical framework based on Pareto optimality predicts that the optimal performance of a multitasker cell that faces trade-offs (*e*.*g*., due to finite resources (Sabi and Tuller 2019; Shoval et al. 2012)) is achieved when its expression is bounded inside a polytope whose vertices are expression profiles optimal at each task, called archetypes (Shoval et al. 2012; Korem et al. 2015; Hart et al. 2015; Hausser et al. 2019) (Figure 1B). The Pareto optimality theory was recently extended to consider an ensemble of cells that are working as a collective to perform the tissue’s tasks (Adler et al. 2019). In the case of collective performance, theory predicts that cells form clusters in gene expression space where they either all concentrate at the polytope’s archetypes (specialists), or all assume an identical composition of task allocation (generalists) (Figure 1B).

**Figure 1:**
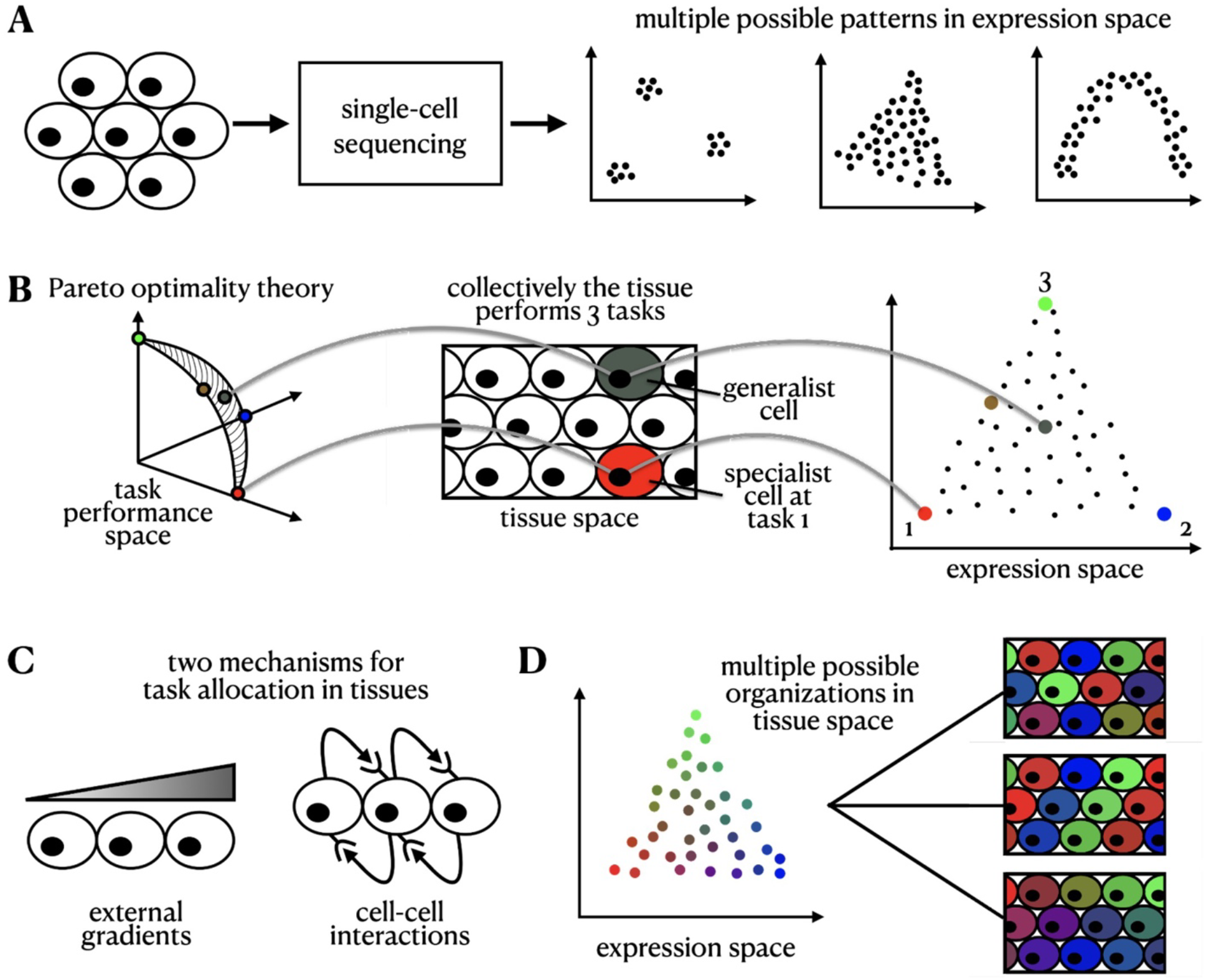
Pareto optimality framework. **(A)** Single-cell RNA-sequencing data in expression space can reveal various low-dimensional structured patterns. **(B)** Mapping of the representation of individual cells across task performance, tissue, and expression space. **(C)** We consider two underlying mechanisms driving the allocation of tasks in the tissue: external signaling gradients and cell-cell interactions. **(D)** Mapping patterns in tissue space with those in expression space is an ill-posed inverse problem -- there are multiple possible tissue compositions consistent with a given expression configuration.

Task performance of cells in a tissue can be affected by exogenous, instructive factors that constrain cellular performance and cannot be altered by direct feedback from the cells such as oxygen and nutrient levels or certain physical barriers. In the case of instructive spatial gradients, a continuum of expression is generated inside the polytope, such that the specialist cells optimal for each task are located in distinct positions in the tissue, where conditions are best for their performance (Adler et al. 2019). For example, variation in expression between individual hepatocytes in the liver lobule (Halpern et al. 2017) and intestinal enterocytes (Moor et al. 2018) can be explained by their spatial positioning in the tissue. However, the variability in expression between cells of other types within the same tissues, for example between fibroblasts in the developing mouse embryo, cannot be easily explained by their spatial position at the tissue scale (Srivatsan et al. 2021).

In addition to instructive constraints, cell-cell interactions, including contact-dependent and secreted signals, also regulate gene expression and coordinate cell performance (Armingol et al. 2021; Belardi et al. 2020) (Figure 1C), such that cells self-organize and use feedback interactions to regulate each other’s performance. This is well-established in cell differentiation: for example, during development, the contact-dependent Delta-Notch pathway controls early neural versus epidermal cell-fate decisions (Beatus and Lendahl 1998). Morphogens, including the Bmp, Wnt, Hedgehog and FGF shape cells’ expression programs and differentiation into distinct cellular fates (Perrimon, Pitsouli, and Shilo 2012; Hogan 1999). In the tissue response to injury, secreted growth factors such as TGF-*β* drive fibroblasts to transition into myofibroblasts, which in turn regulate the wound healing process (Gabbiani 2003; Baum and Duffy 2011). Immune cells constantly sense the landscape of cytokines and chemokines produced by other cells to determine the nature of their response (Altan-Bonnet and Mukherjee 2019).

While cell-cell communication is a key mechanism that governs cell differentiation, how cell-cell communication promotes division of labor within a given cell type remains unclear. Could a continuum within a polytope in gene expression space emerge when cells co-organize to control their task allocation? What would be the physical arrangement of cells in the tissue when their task allocation is controlled by cell-cell interactions locally or at longer ranges? Can we leverage the Pareto optimality framework to predict the spatial arrangements of cells in the tissue based on their gene expression profile (Figure 1D)?

Here, we develop and apply a theory that considers the optimal trade-off between tasks in the tissue when cells communicate their specializations to each other. Unlike instructive factors, where the conditions of performing tissue-wide tasks are guided extrinsically, in this framework, the cells use feedback interactions and self-organize to divide labor in an optimal way. The theory predicts a diversity of patterns both in tissue (physical) and gene expression spaces that emerge from different types of cell-cell interactions that are distinct from patterns that emerge from instructive signals. We apply the theory to single-cell RNA-seq (scRNA-seq) and spatial transcriptomics data by comparing the spatial organization and expression patterns of mouse colon fibroblasts and intestinal mature enterocytes and by constructing ligand-receptor crosstalk networks across specialist cells from single-cell gene expression data of fibroblasts and macrophages. This framework can provide insights into whether expression patterns originate from instructive constraints or cell-cell interactions in diverse biological systems.

## Results

### Modeling cell-cell communication under the Pareto optimality framework generates a variety of patterns in tissue and gene expression space

To model the Pareto-optimal expression profiles of cells in a tissue, we consider how cells collectively contribute to the tissue by performing several tasks. As was previously presented (Shoval et al. 2012; Adler et al. 2019), we model this trade-off by considering that each task is best performed at an optimal expression profile, *G_t_*\* (or an optimal task allocation) and shows a decline in performance as cells move further away from *G_t_*\* in gene expression space. We define the total performance function of a tissue, *F*, as a product over the performance of all the tasks that need to be collectively performed by the cells in the tissue, summing over the contribution of each cell to the performance in each task (Adler et al. 2019) (Methods).

To model the effect of cell-cell interactions on optimal task allocation, we introduce an interaction term, *H*_*t*_, which captures how a cell’s performance is influenced by the performance of its neighboring cells. We explore the effect of varying the range of the interaction by varying the size of the neighborhood of each cell (*N*_*c*_). The contribution of each cell (*c*) in task *t* is therefore the product of two components; a *self-component, Pt*, which is a function of cell *c*’s gene expression profile (*G*_*c*_), and an *interaction component, H*_*t*_, which is a function of the average *P*_*t*_ of the neighboring cells of cell *c* (Figure 2A).

**Figure 2:**
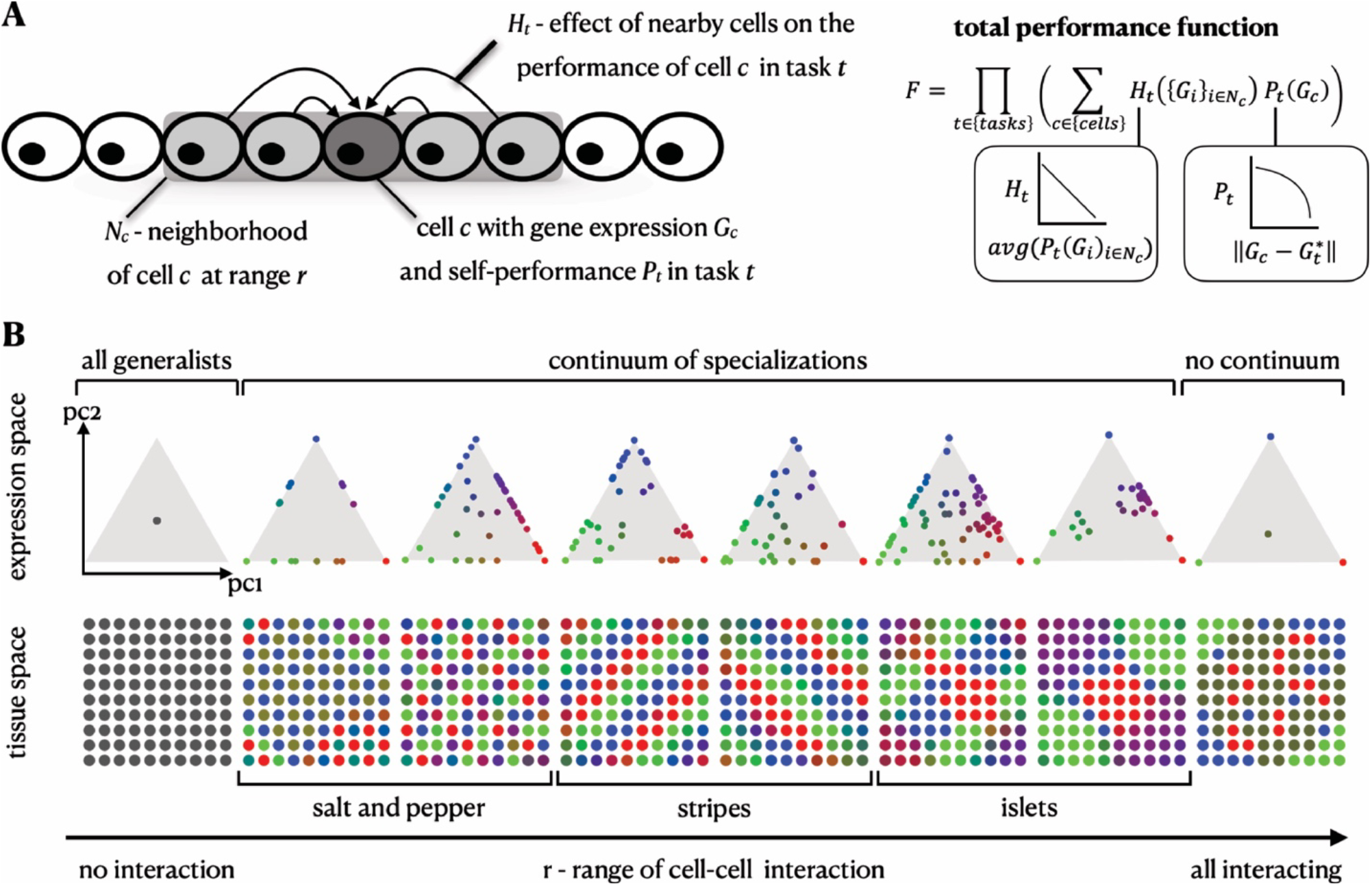
A variety of tissue and expression patterns emerge from the Pareto optimality framework with a cell-cell communication mechanism of lateral inhibition. **(A)** Theoretical framework of Pareto optimality with cell-cell interactions - Cell *c*’s performance is affected by the performance of its neighbor cells in the tissue, *N*_*c*_. The total performance function, *F*, is a product over the tasks, where the performance in each task is the sum over the contribution of all cells considering a self-component (*P*_*t*_) and the effect of interaction with nearby cells (*H*_*t*_). **(B)** Simulation results of lateral inhibition in expression and tissue space. Varying the range of cellular interactions produces diverse tissue patterns akin to patterns observed in real tissues.

The interaction term, *H*_*t*_, can generally represent different types of interactions, including positive and negative effects on performing the same task. Here, we focus on lateral inhibition, where a cell’s performance in task *t* declines if its neighboring cells exhibit high performance in the same task (Figure 2A, Methods). We consider a representative example of a 2D spatial grid of 100 cells that need to perform three tasks and compute the expression profiles (or task allocations) of the cells that maximize *F* (Methods). We discuss tissue dimensions, number of tasks, and other types of interactions in the supplementary information (SI) (Figure S1-2).

In the absence of cell-cell interactions, the tissue reaches optimal performance when all cells are generalists, equally performing all tasks, but when interactions are present, the cells span multiple qualitatively distinct arrangements within the Pareto front in gene expression space (a triangle in the case of three tasks), and a diversity of spatial patterns in the tissue, depending on the range of interactions (Figure 2B). Specifically, short-range / direct interactions with nearest neighbors lead to the formation of a salt-and-pepper pattern in tissue space, where the cells are confined to the circumference of the polytope in gene expression space. Increasing the range of lateral inhibition interactions drives the cells to fill the polytope with a continuum of profiles spanning both specialist and generalist cells. At the other extreme, when all cells are mutually interacting irrespective of location, the optimal solution partitions the cells to specialists and generalists (no continuum). In tissue space, the model gives rise to a range of physical patterns often observed in tissues in development and homeostasis (Gur et al. 2020), including stripes and islets of the different specialist cells (Figure 2B).

### Both instructive gradients and cell-cell interactions lead to continua of gene expression profiles but distinct spatial patterns

When simulating lateral inhibition of cell-cell interactions, a continuum of gene expression profiles can emerge (Figure 2B), similarly to the continuum generated by external gradients (Figure 3A-E, Methods) (Adler et al. 2019), but with a key difference in the spatial configuration of the cells. With instructive monotonic gradients, cells that are physically proximal within the tissue can exhibit many times, on average, similar expression profiles (Nitzan et al. 2019) and thus similar task allocation. In this case, specialist cells of distinct specializations are expected to be located far from each other within the tissue, and similarity in gene expression and physical proximity are congruent. Conversely, in a population of cells whose expression profiles are driven by lateral inhibition, different specialist cells can be close to each other in tissue space (Figure 3D, E), such that gene expression and tissue space can be incongruent. We demonstrate this quantitatively by comparing the distribution of pairwise distances of cells in tissue space (“physical distance”) and in task allocation space (“task distance”), where a cell’s task allocation is the normalized gene expression distance from each of the archetypes (Figure 3C, F). We then construct a null model for assessing the statistical significance of the physical locations of cells using random permutations (Methods). Simulations with instructive monotonic gradients show a statistically significant Pearson correlation between pairwise distances in gene expression and physical spaces (Figure 3C, F, Methods). When a combination of both instructive gradients and lateral inhibition governs the simulation, the Pearson correlation of physical and gene expression pairwise distances generally increases as the strength of the effect of the external signaling gradients increases (Figure S3).

**Figure 3:**
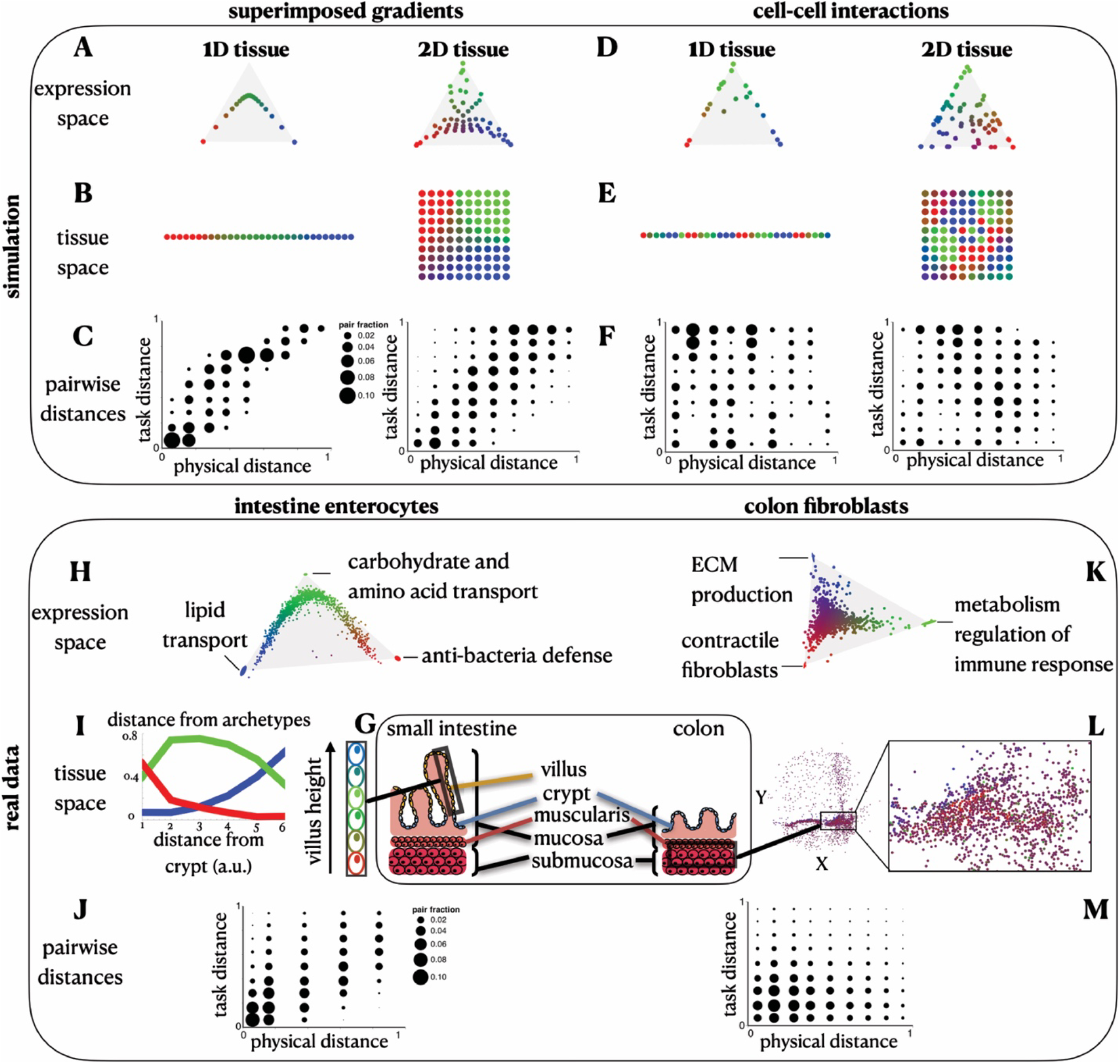
Distinct spatial patterns emerge from external gradients and cell-cell interactions. **(A-B)** The Pareto-optimal solution of cells that are affected by an external gradient across a 1D and a 2D tissue in gene expression **(A)** and tissue **(B)** space. **(C)** With external gradients, the pairwise expression distances (y-axis) versus the pairwise physical distances (x-axis) show high Pearson correlation (for 1D: corr=0.88, p-val<0.001, for 2D: corr=0.72, p-val<0.001). **(D-E)** The Pareto-optimal solution of cells that are affected by cell-cell interactions in a 1D and a 2D tissue in gene expression **(D)** and tissue **(E)** space. **(F)** With cell-cell interactions, the pairwise expression distances (y-axis) versus the pairwise physical distances (x-axis) are anticorrelated (in terms of Pearson correlation, for 1D, corr=-0.1, p-val=0.002, for 2D, corr=-0.13, p-val<0.001). **(G)** Schematics of the colon tissue including the intestinal villi and the muscularis layer. **(H)** Gene expression profile of enterocytes where the cells are colored by their task specializations (combinations of red, green and blue colors representing the three tasks). **(I)** Arrangements of enterocytes along the crypt-to-villus axis. The distance of the cells from each archetype is plotted as a function of the distance from the crypt. **(J)** The pairwise distances in expression versus physical space of enterocytes show high Pearson correlation (corr=0.67, p-val<0.001). **(K)** Gene expression profile of colon fibroblasts from Slide-seq data (Avraham-Davidi et al., 2022), where the cells are colored by their task specializations. **(L)** Spatial arrangements of the fibroblasts in the colon tissue. **(M)** The pairwise distances in expression versus physical space of fibroblasts show negative Pearson correlation (corr=-0.1, p-val<0.001).

### Both intestinal enterocyte and colon fibroblast distributions show task specialization, but only enterocytes’ task allocation can be explained by a global instructive gradient

To test our predictions in the context of real biological systems, we turned to two datasets, to infer whether the distribution of gene expression profiles is dominated by either instructive gradients or cell-cell interactions. The first system consists of mature enterocytes from the villus of the mouse small intestine, dissected into six regions using laser capture microdissection and then dissociated and profiled by scRNA-seq (Moor et al. 2018). Multiple nutrients and resource gradients were previously suggested to influence the expression profiles of enterocytes along the crypt-to-villus axis (Figure 3G, I). The second system consists of fibroblasts in the mouse colon assayed *in situ* using the spatial transcriptomics method Slide-seq (Avraham-Davidi et al., 2022) (Figure 3G, L).

Next, we test whether expression profiles of colon fibroblasts are governed by an instructive gradient, similar to enterocytes in the small intestine, and what is the role of cell-cell interactions in regulating their heterogeneity at steady state. As expected, enterocyte profiles follow a one-dimensional continuum of expression bounded within a triangle in gene expression space, as previously shown (Adler et al. 2019), consistent with a trade-off in enterocytes between three tasks: lipid transport, carbohydrate and amino acid transport, and antibacterial defense (Figure 3H). Specialist enterocytes are found in distinct positions in the tip, middle and bottom of the villus (Methods). (This pattern was observed among mature enterocytes that line the villus from the small intestine; enterocytes from other regions may show distinct patterns.) Conversely, Slide-seq beads capturing fibroblasts span an expression continuum within a triangle (p-val<10^−4^, based on the ParTI algorithm (Hart et al. 2015)). The genes enriched near each of the fibroblast archetypes suggest task specialization in extracellular matrix (ECM) production, contractile functions, and metabolism and regulation of immune response (Figure 3K, Methods).

Unlike the gradual change in enterocyte profiles along the crypt-to-villus axis (Figure 3I), the distinct colon fibroblast archetypes are often close to each other in the tissue space (Figure 3L). Computing the expression and physical pairwise distances for both cell type populations, we find a significant Pearson correlation for enterocytes, in line with the assumption of a dominant monotonic gradient along the villus axis, while the signature for fibroblasts is not consistent with monotonic spatial gradients alone (Figure 3J, M, Methods). This distinction is observed even when we simulate in the Slide-seq data the coarse laser cutting dissection approach that was applied for enterocytes in the small intestine. This shows that the difference in correlation between physical and task distances observed for mature intestinal enterocytes and colon fibroblasts is not merely explained by the difference in experimental assays (Figure S5A-B). Although colon fibroblasts are known to be influenced by morphogenic signals such as Wnt produced by deep-crypt secretory cells (Shoshkes-Carmel et al. 2018; Sasaki et al. 2016; Gehart and Clevers 2019), the spatial mixture of fibroblast archetypes within the crypt suggests that fibroblasts are also influenced by additional mechanisms (Figure S5C). We next turn to examine whether cell-cell interactions between fibroblasts play a central role in regulating fibroblast heterogeneity.

### An archetype crosstalk network for colon fibroblasts highlights specific ligand-receptor interactions as potential mechanisms for optimal task allocation

To explore whether cell-cell interactions between fibroblasts play a role in fibroblast task specialization, we developed a method to construct crosstalk networks between archetypes of different specializations based on ligand-receptor interactions (Figure 4A, Methods). Specifically, we construct a directed graph where vertices are the different archetypes, and a directed edge connects from archetype A to archetype B if a ligand whose expression is enriched towards archetype A in gene expression space has a corresponding receptor whose expression is enriched towards archetype B (Figure 4A, Methods). Here we consider interactions within the same cell type population (*e*.*g*., fibroblasts) and their effect on task performance, although task performance of fibroblasts may be additionally affected by interaction with other cell types, which we do not consider.

**Figure 4:**
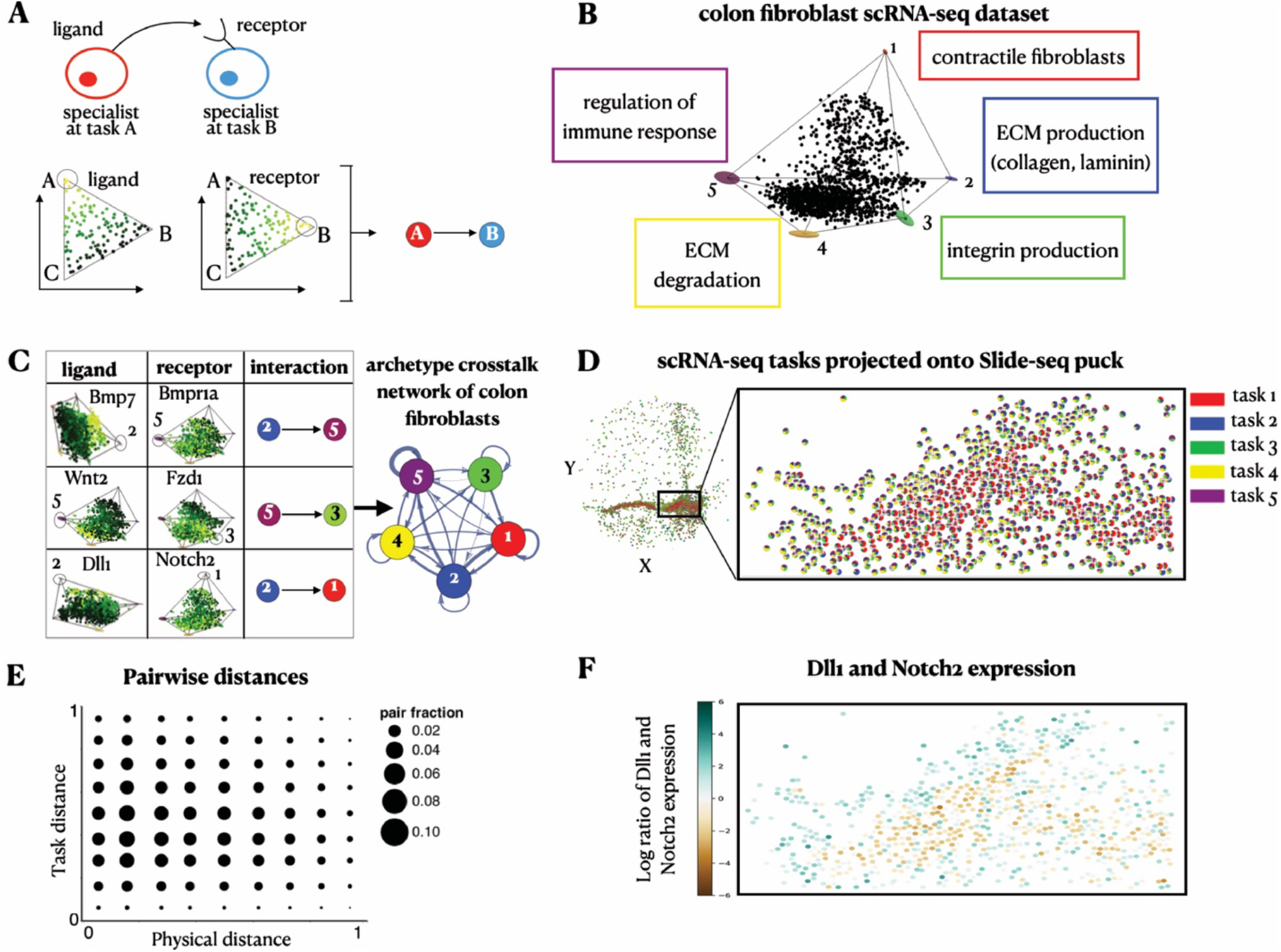
Inferring archetype crosstalk networks of colon fibroblasts based on ligand-receptor interactions. **(A)** Interactions between specialist cells are inferred from enrichments of ligands and their corresponding receptors. We use a directed edge to represent a pair of a ligand and its corresponding receptor that are enriched near each of the archetypes it connects. **(B)** A projection of gene expression profiles of scRNA-seq colon fibroblasts (Muhl et al. 2020) on the first three PCs. Fibroblasts fill in a 5-vertex polytope (p-val<10^−4^). **(C)** A table showing examples of ligands and their respective receptors that are enriched near the archetypes. We plot the complete archetype crosstalk network inferred for the colon fibroblasts where the thickness of each edge corresponds to the number of ligand-receptor pairs enriched between its vertex archetypes. **(D, E)** Using TACCO (Mages et al., 2022), we compute a mapping from fibroblast cells assayed by scRNA-seq and Slide-seq beads based on their expression agreement. Using the mapping, we **(D)** infer the scRNA-seq task components for each bead (depicted in pie-charts per bead) and **(E)** compute the corresponding correlation of pairwise task distances versus physical distances (corr=-0.04, p-val < 10^-4). **(F)** Projection of the scRNA-seq expression profiles onto the Slide-seq beads. To view the expression of Delta and Notch, we image their log-ratio. Beads whose inferred Delta expression is greater/less than their Notch expression lean towards a turquoise/brown shade, respectively.

We first applied this approach to scRNA-seq data of mouse colon fibroblasts (Muhl et al. 2020) (which more sensitively detects receptor and ligand gene expression than Slide-seq). Colon fibroblast profiles span a 5-vertex expression polytope with five archetypes corresponding to fibroblast contractile functions, ECM production including collagen and laminin, integrin production, ECM degradation, and regulation of immune response (Figure 4B, Methods). These archetypes are in line with fibroblast categories that were recently described (Buechler et al. 2021). Interestingly, the pareto optimality analysis reveals two distinct archetypes of collagen production and contractile functions, suggesting that there are two subpopulations performing myofibroblast functions.

The constructed archetype crosstalk network based on a database of ligand-receptor pairs (Shao et al. 2021) shows that all colon fibroblast archetypes potentially interact with each other (Figure 4C) (unlike enterocytes, discussed below). These interactions include secreted signaling molecules, such as Bmp, Wnt and Fgf, which are known to be produced by fibroblasts in the colon and play a significant role in regulating tissue organization (Roulis and Flavell 2016) (Figure 4C). Interestingly, the ECM production and contractile fibroblast archetypes are enriched for Delta (Dll1) and Notch (Notch2) respectively, suggesting that fibroblasts may use contact-dependent signals and lateral inhibition to regulate their task specializations.

Next, we mapped the scRNA-seq data to Slide-seq data using TACCO (Methods, (Mages et al., 2022)), first by annotating Slide-Seq beads with ‘fractions’ of cell type annotations, and then projecting the scRNA-seq task annotations and expression profiles onto tissue space (Figure 4D, Figure S6). Like the Slide-seq task annotations (Figure 3L), the scRNA-seq task annotations also present a clear boundary between the contractile fibroblast (red) and ECM-related (blue and yellow) archetypes and lack global smoothness of task annotations (Figure 4E, corr=-0.04, p-val<10^−4^), in support of mechanisms beyond spatial smooth signals controlling the behavior of cells. Notably, immune response fibroblasts are interspersed with contractile fibroblasts in space, but clearly separable as archetype (Figure 4E). The projected expression of strong ligand-receptor interactions inferred from the scRNA-seq, such as the Dll1-Notch2 pair enriched in ECM production and contractile fibroblast archetypes, respectively, along the spatial boundary of these archetypes suggests the involvement of cellular interactions in setting the task allocations across the tissue (Figure 4F). The spatial proximity between ECM producing cells and contractile specialists may be important where performance of contractile functions may depend on the ability of the cells to attach to matrix factors produced by the ECM producing cells. This is supported by the fact that contractile fibroblasts show enrichment of integrin (Itgb1) which serves as a receptor for collagen (Col1a1, Col1a2, Col4a1, Col4a5, Col4a16 and laminin (Lama5, Lamb1) produced by the ECM producing cells. Additionally, the ECM producing cells are enriched with expression of thrombospondin (Thbs1) – an adhesive glycoprotein that mediates cell-to-cell and cell-to-matrix interactions that can bind to integrin.

### Archetype crosstalk networks can be used to dissect communication-driven tissue organization from non-spatial single-cell profiles

Even if positional information of the cells is not available, archetype crosstalk networks can highlight potential cell-cell interactions that regulate cell specialization. We demonstrate this for fibroblast and macrophage profiles from human lungs (data from (Adams et al. 2020)). Lung fibroblasts span five archetypes that are overall similar to the five colon fibroblast archetypes above, with fibroblast contractile functions, ECM production, ECM degradation and immune response regulation as distinct archetypes in both tissues. Specifically, lung fibroblasts specialize in (1) regulation of immune response (with enriched cytokines (CCL2), chemokines (CXCL2, CXCL3, CXCL8) and interleukins (IL32, IL33)); (2) ECM degradation (ADAMTS1, ADAMTS4, MMP14); (3) protein biosynthesis and metabolism, including collagen production; (4) ECM production and regulation; and (5) contractile fibroblast functions (Figure 5A, Methods). Lung macrophages fill in a tetrahedron in expression space and trade-off between ECM degradation, phagocytosis, metabolism of lipids, proteins, glucose and fatty acids, and pro-inflammatory response (Figure 5C, Methods).

**Figure 5:**
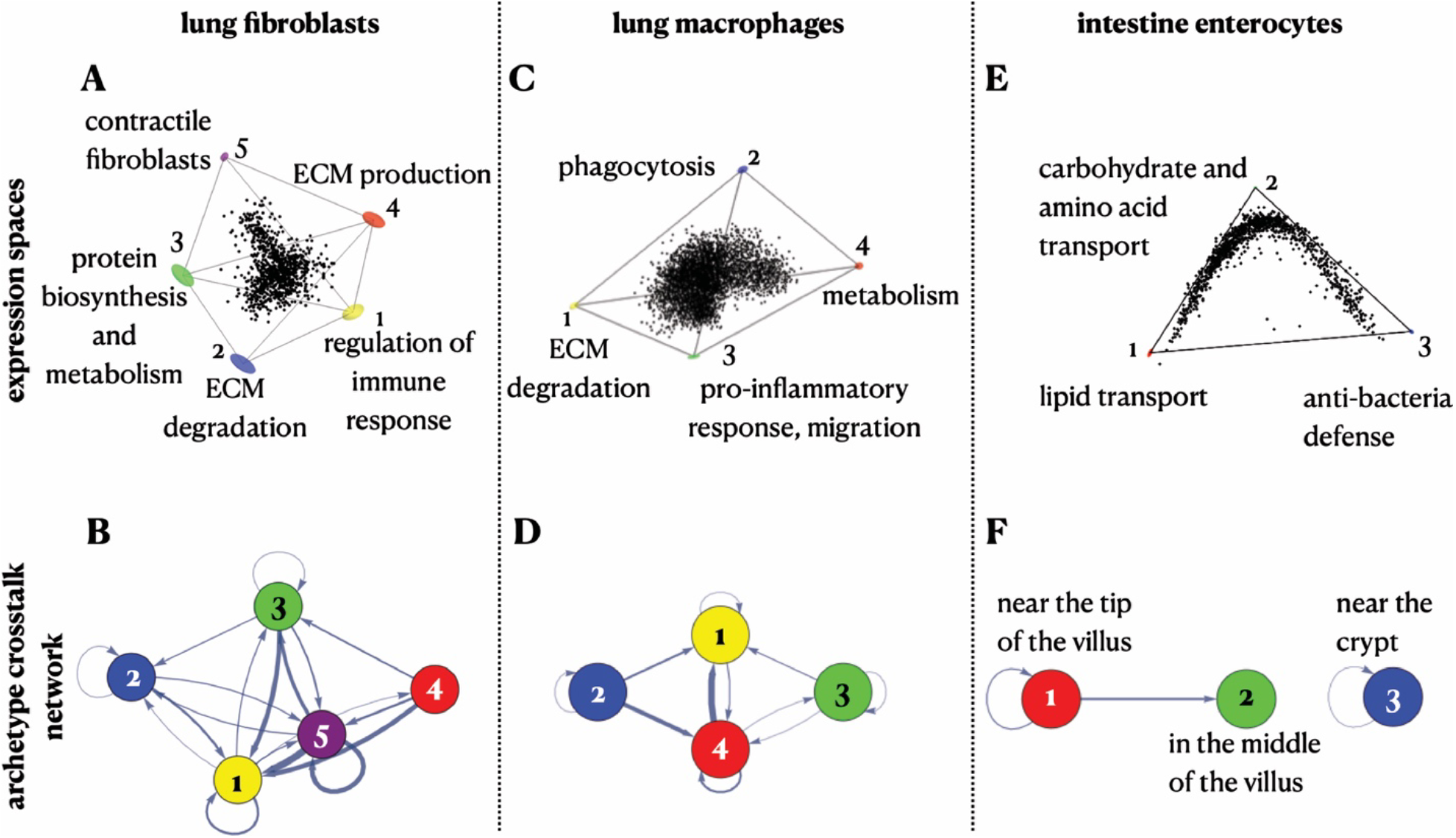
Archetype crosstalk networks can help estimate the role of cell-cell interactions in shaping task allocation even without spatial information. **(A)** Human lung fibroblasts (from (Adams et al. 2020)) fit in a 5-vertex polytope in expression space (p-val=0.004) and show 5 archetypes that correspond to regulation of immune response, ECM degradation, protein biosynthesis and metabolism, ECM production and myofibroblasts specializations. **(B)** Archetype crosstalk network of lung fibroblasts. **(C)** Human lung macrophages (from (Adams et al. 2020)) fit in a tetrahedron in expression space (p-val<10^-4) and show 4 archetypes that correspond to ECM degradation, phagocytosis, pro-inflammatory response and metabolism specializations. **(D)** Archetype crosstalk network of lung macrophages. **(E)** Expression profile of intestine enterocytes (from (Moor et al. 2018)). **(F)** Archetype crosstalk network of intestine enterocytes.

The archetype crosstalk networks for lung fibroblasts and macrophages show that both cell types exhibit strong connectivity between the different archetypes within each cell type population (Figure 5B, D) (See Methods for a comparison between the connectivity in the archetype crosstalk networks to connectivity in networks of non-specialist cells, Figure S7). In contrast, in the intestinal enterocytes’ archetype crosstalk network, not all archetypes interact with each other, and the interactions are restricted to spatially proximal cells (Figure 5E-F, Methods). Thus, even without spatial data, a pattern of strong connectivity between the different archetypes within a given cell type population suggests that cell-cell interactions play an important role in regulating the division of labor between the cells.

## Discussion

Cells work together in tissues and contribute to their collective function in homeostasis and their dysfunction in disease. In this work, we studied how cell-cell communication shapes the distribution of cellular gene expression. Through simulations based on our theoretical framework and its application to real data, we conclude that self-organization mechanisms can explain a diversity of patterns in both task and tissue space. Additionally, we construct a framework for distinguishing the effects of cell-cell communication and instructive gradients by mapping ligand-receptor interactions between specialized cells.

Inferring the underlying mechanism for division of labor is a challenge since multiple potential mechanisms may result in the same phenotype. As comprehensive spatial transcriptomics data become more prevalent and extended to capture three-dimensional tissue structure (Wang et al. 2018) and causation (Legnini et al. 2022), our framework can be used to better infer or narrow down the potential underlying mechanisms of collective cellular division of labor.

Our theory currently focuses on the principles underpinning the optimal division of labor of cells within a given cell type population. Future work accounting for multiple different cell type populations could depict a more holistic image of tissues, which would enable, for example, to explore how crosstalk between fibroblasts, epithelial, and immune cells in the colon regulates their heterogeneity and division of labor.

In this work, we focused on characterizing the division of labor among cells in a steady state of a developed tissue. Expanding the theoretical framework to consider Pareto optimality in a dynamical setting can provide insights into developmental strategies and their regulation. Another question that can be explored using the Pareto framework applied to time-resolved data is whether cells converge to specific specializations or can the same cell switch between different specializations when needed, for example in the context of epithelial mesenchymal transition as was recently suggested (Cook and Wrana 2022). Finally, our framework can be used to explore the transition of tissues to pathological states, and particularly, to study how the role of cell interactions in regulating fibroblast heterogeneity is reshaped in the context of fibrosis and cancer.

## Acknowledgments

We thank Uri Alon, Johanna Klughammer and members of our labs for meaningful discussions. This work was supported by the EMBO Long-Term Fellowship (ALTF 304-2019), the Zuckerman STEM Leadership program, and the Israel National Postdoctoral Award Program for Advancing Women in Science (MA), the Center for Interdisciplinary Data Science Research at the Hebrew University of Jerusalem (NM and MN), a DFG research fellowship (MA 9108/1-1) (SM), NIH grants R01HL127349, U01HL145567, U01HL122626, and the Three Lakes Foundation (NK), the BroadIgnite philanthropic grant (AG), the Howard Hughes Medical Institute (AR and RM), the Klarman Cell Observatory (AR), the Blavatnik Family Foundation (RM), a grant from the NIH AI144152 (RM), Azrieli Foundation Early Career Faculty Fellowship (MN), Horizon Europe ERC Grant 101040660 (MN), and an ISF Research Grant 1079/21 (MN).

## Author Contributions

Conceptualization - MA, NM, AG, AR, RM, and MN; Methodology - MA, NM, AG, and MN; Software - MA, NM, SM, and MN; Validation - MA, NM, and AR; Formal Analysis - MA, NM, and AG; Investigation - IAD, TSA, and NK; Resources - IAD, TSA, NK, AR, and MN; Writing – Original Draft - MA, NM, AG, AR, RM, and MN; Writing – Review & Editing - All Authors; Visualization - MA, NM, and AG; Supervision - EZM, AR, RM, and MN.

## Methods

### Cell-cell communication model under the Pareto optimality framework

The Pareto optimality framework introduced in (Adler et al. 2019) (set to allocate cells’ gene expression towards a maximal, collective performance of the tissue) is formalized by the constrained maximization of the total performance function, F:

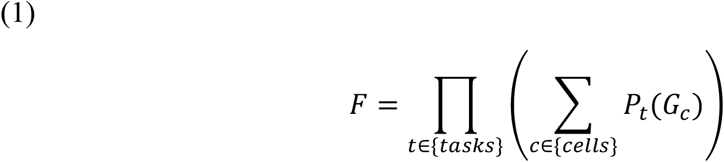

Where, *G*_*c*_ is a vector of cell c’s gene expression, that is constrained to lie in the polytope whose vertices are the archetypes’ gene expression, that is 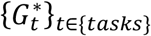. The performance of a cell for task t, *P*_*t*_(*G*_*c*_), is computed as 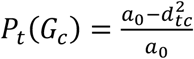 where 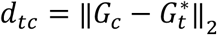. (Euclidean distance of cell c from the archetype t) and *a*_0_ is a constant introduced to avoid negative performance (default:*a*_0_ = 2). Summation over the cells describes the equal, linear contribution of each cell’s performance in task t to the collective performance of task t. The product over the performance in each task expresses the need to excel in all tasks simultaneously.

To model the effect of a cell’s local environment, we consider cells arranged in an acyclic 1D or 2D grid. *N*_*c*_, the set of cell c’s neighbors is set to be cells within order r (range) from c.

For interactions of lateral inhibition of the same task, we factor in 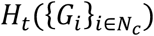, a diminishing factor (≤ 1) introduced by the performance in task t of cell c’s neighbors:

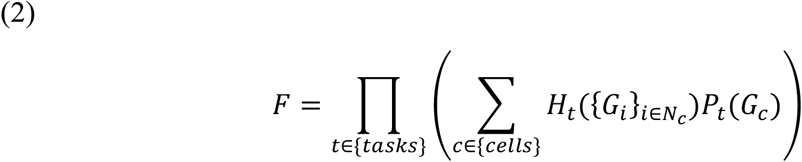

For simplicity, we set *H*_*t*_ to be: 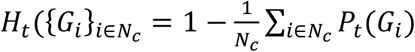

We implemented the optimization problem both in Python and Mathematica. Examples, varying tissue dimensions, and number of tasks, are depicted in Figure S1.

### Expression and physical distances are significantly correlated under instructive gradients, but not for interactions

We wish to compare the expression and tissue space optimized with external gradients or with interactions. For external gradients, we use the setup previously proposed in (Adler et al. 2019) where:

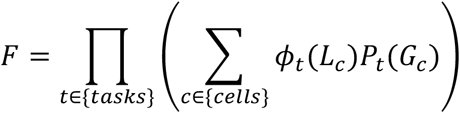

Where *L*_*c*_ is cell c’s location within the tissue. *ϕ*_*t*_(*L*_*c*_) is a coefficient defined by the external gradient that weighs cell c’s performance according to gradients at its location (*L*_*c*_) in the tissue. Exact gradients set for 1D, and 2D tissues are *ϕ*_1_(*x*) = *x*, *ϕ*_2._(*x*) = 1 − *x*, and *ϕ*_1_(*x*, *y*) = 1 −*x*, *ϕ*_2._(*x*, *y*) = 1 − *y*, *ϕ*_3_(*x*, *y*) = 1, respectively, and we use 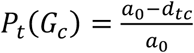for evaluating the performance in correspondence to the simulation in (Adler et al. 2019) (Figure 3A-B).

To plot binned expression-vs-physical-distances, we bin (separately) cell-pairs’ task allocation distances and cell-pairs’ physical locations: (1) compute cell-to-cell Euclidean distance (expression or physical distances), (2) remove extreme distances (> 99 percentile), (3) normalize(divide by max distance) to contain in 0-1 range, (4) bin into equal-range all distances (using 10 bins, for enterocytes, physical distances collapse into 5 bins due to low spatial resolution).

Then for each (expression distance bin x physical distance bin), we plot a point of size corresponding to the fraction of cell pairs (number of pairs / total number of pairs) falling within this set of bins. Correlation values are computed between the plain Euclidean expression/physical distances and compared to null correlation values (1000 repeats) that are generated by permuting cells’ locations.

### Analysis of single-cell RNA-seq data of intestinal enterocytes

We re-analyzed scRNA-seq data of mouse intestinal enterocytes (data from (Moor et al. 2018)) where we preprocessed and normalized the data and considered the triangle that encloses the data as was previously described in (Adler et al. 2019). See Table S1 for a full list of enriched genes near the enterocyte archetypes.

To plot the pairwise distances in expression and tissue space, we consider every pair of enterocytes in the data. For distance in expression space, we compute the Euclidean distance in task space between every pair of cells. For distance in spatial space, we consider the spatial height of the cells along the crypt-to-villus axis that was inferred by Moor et al. (Moor et al. 2018), and compute the difference between the heights of every pair of cells.

The full list of enriched ligand-receptor pairs in the enterocytes data that were used to plot the archetype crosstalk network in Figure 5F can be found in Table S2.

### Analysis of Slide-seq data of colon fibroblasts

We analyzed Slide-seq data of mouse colon fibroblasts from Avraham-Davidi et al. (Avraham-Davidi et al., 2022), where we considered fibroblasts as beads that received a fibroblast score higher than 0.5 (Mages et al., 2022). In the main text we focus on a specific slide from the dataset (puck #20), but we analyzed other slides included in the data and found similar results (see Figure S4A-D for results of another puck in the data). Puck #20 includes 2559 fibroblasts that show expression in 13,520 genes. To preprocess the expression data, we used ‘Sanity’ – a recently developed method to normalize single-cell data and to infer the transcriptional activity of genes (Breda, Zavolan, and van Nimwegen 2021). Following the Sanity normalization, we removed genes with low expression (log_10_(average expression) < -11) and low expression variance (standard deviation < 0.1) across beads, which left us with 1047 genes. We note that these thresholds for selection of genes does not affect the distribution of expression profiles in expression space and analyzing all 13,520 genes yields similar results.

We next used the ParTI package in Matlab to fit the fibroblast expression data to a polytope (Hart et al. 2015). We find that fibroblasts fit well within a triangle (p-val<10^,4^, we also tested for 4 archetypes which yield a good fit with p-val=0.001). We used ParTI to find genes that are enriched near the archetypes to infer the tasks the archetypes are specializing in. The enriched genes near the first myofibroblast archetype are: Malat1, Actg2, Myh11, Tpm1, Pdlim3, Acta2, Mylk, Cnn1, Smtn, Tagln, Des, Actb, mt-Nd4, Flna, Myl9. The enriched genes near the second ECM production archetype are: Col3a1, Col1a2, Gsn, Dcn, Sparc, Dpt. The enriched genes near the third metabolism/immune response archetype are: Lars2, Cmss1, Hexb, Camk1d, Tpm2. We find similar enriched genes in other pucks in the dataset (Figure S4D).

To plot the pairwise distances in expression and tissue space, we consider every pair of fibroblasts in the data. For distance in expression space, we compute the Euclidean distance in task space between every pair of cells. For distance in spatial space, we consider the spatial (x, y) coordinates of the cells from the slide-seq data and compute the Euclidean spatial distance between every pair of cells.

### Construction of archetype crosstalk networks based on ligand-receptor interactions

To construct archetype crosstalk networks from single-cell expression data, we first fit the data to a polytope. Once the polytope and archetypes are defined, we consider the genes that are enriched near the archetypes and use available ligand-receptor pairs datasets (we used CellTalkDB: (Shao et al. 2021)) to search for enriched ligand-receptor pairs. We used the package IGraphM in Mathematica 12.1.1.0 and the functions Graph, IGEdgeMap, IGEdgeProp to build a weighted directed graph where a directed edge from archetype A to archetype B represent an enriched ligand near Archetype A and its corresponding receptor enriched near archetype B. The weights on the edges correspond to the number of ligand-receptor pairs that link the two archetypes.

### Analysis of single-cell RNA-seq data of colon fibroblasts

We analyzed scRNA-seq data of mouse colon fibroblasts from Muhl et al., 2020. The data includes 1646 cells that show expression in 30,920 genes. Following the Sanity normalization, we removed genes with low expression (log_10_(average expression) < -14) and low expression variance (standard deviation < 0.1) across single cells, which left us with 8479 genes. We next used the ParTI package in Matlab to fit the fibroblast expression data to a polytope. We find that fibroblasts fit well within a 5-vertex polytope (p-val<10^,4^, we also tested for 3 and 4 archetypes which yield a good fit with p-val<10^,4^). We find hundreds of enriched genes near the 5 fibroblast archetypes (see Table S3 for the full table of enriched genes). The full list of enriched ligand-receptor pairs in the scRNA-seq colon fibroblasts data that were used to plot the archetype crosstalk network in Figure 4E can be found in Table S4.

### Mapping single-cell data onto Slide-seq positions of colon fibroblasts

To further support the role of cell-cell interactions in guiding the spatial task allocations for colon fibroblasts, we map the high-quality single-cell data onto the fibroblast-dominated (probability fibroblast > 0.5) Slide-seq beads. To this end, we deploy TACCO, mapping the Slide-seq data onto single-cell data as a reference (Mages et al., 2022). We use TACCO’s platform normalization booster to overcome platform biases and Optimal Transport to map solely based on gene expression similarity to the reference expression profiles. Using the cell-to-bead mapping, we compute the single-cell task allocation and the inferred expression per bead.

### Analysis of single-cell RNAseq of human lung fibroblasts and macrophages

We analyzed single-cell RNA-seq data of human lung fibroblasts from Adams et al. 2020. The data includes 1051 cells from the control samples. Following the Sanity normalization, we removed genes with low expression (log_10_(average expression) < -11) and low expression variance (standard deviation < 0.05) across single cells, which left us with 6934 genes. We also removed a small fraction of outlier cells that skewed the data that show low average expression (log_10_(average expression) < -9.9), which left us with 945 cells.

The human lung macrophage data from Adams et al. (Adams et al. 2020) includes 25142 cells (macrophages and monocytes). Due to the large number of cells, we randomly sampled 5000 cells to carry on with the analysis. Following the Sanity normalization, we removed genes with low expression (log_10_(average expression) < -11) and low expression variance (standard deviation < 0.05) across single cells, which left us with 6273 genes.

We used the ParTI package in Matlab to fit the fibroblast and macrophage datasets to a polytope. We find that the fibroblast data fits well in a 5-vertex polytope (p-val=0.004 with the Sisal algorithm to find the archetypes, p-val=0.067 with the PCHA algorithm to find the archetypes), and the macrophage data fits well in a tetrahedron (p-val<10^-4 with the Sisal algorithm, p-val=0.067 with the PCHA algorithm). We repeated the enrichment analysis using both PCHA and SISAL algorithm and we found that the enriched genes near the archetypes were very similar using both algorithms suggesting that the results are robust to different methods of finding the archetypes’ positions.

Tables of full lists of enriched genes near the fibroblast and macrophage archetypes can be found in Tables S5, S6. The full list of enriched ligand-receptor pairs in the lung fibroblasts and macrophages datasets that were used to plot the archetype crosstalk network in Figure 5B, D can be found in Tables S7, S8.

## Supplementary information figures

**Figure S1:**
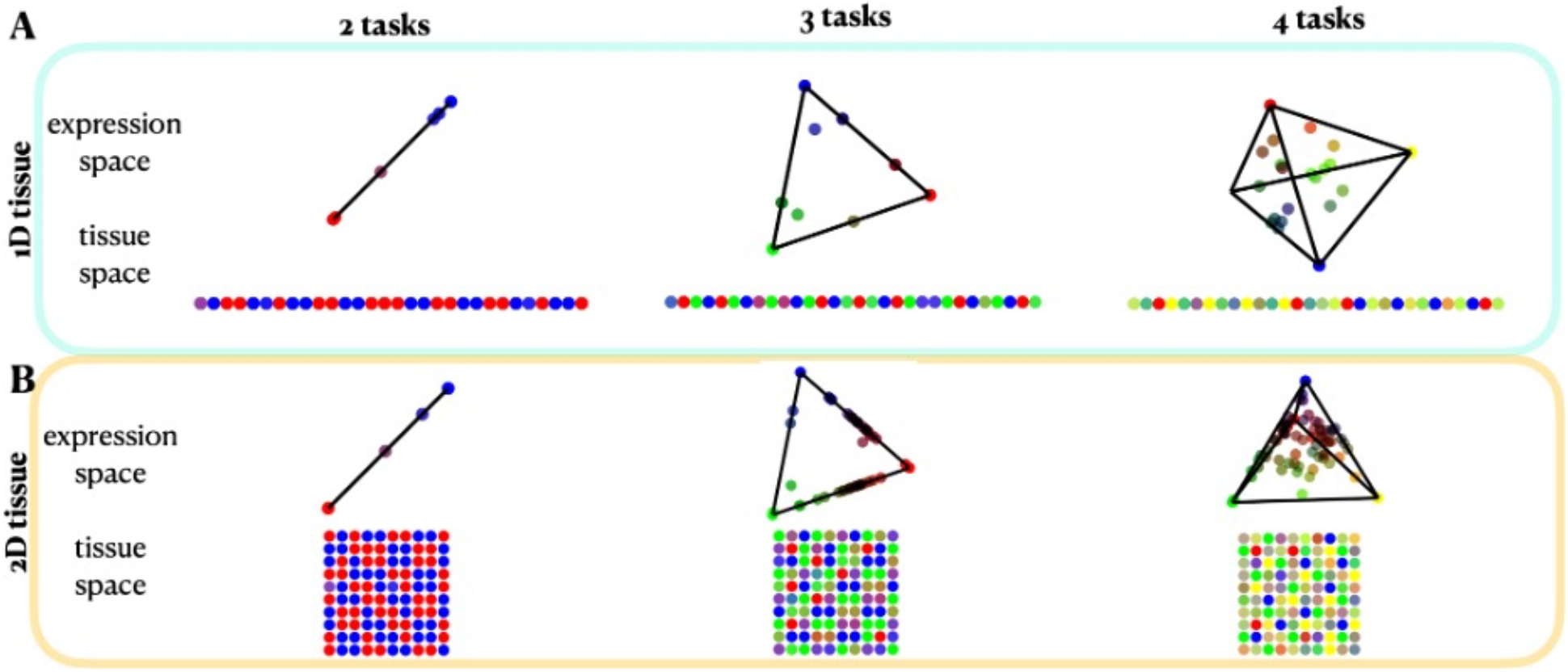
Simulation results with lateral inhibition. Columns describe the number of tasks (2, 3, and 4). Rows describe the dimension of the spatial organizations of cells in the tissue for 1D grid with 30 cells **(A)**, and 2D grid with 100 cells **(B)**. In all cases, a neighborhood of range 2 is considered. We plot the expression space (vertices represent archetypes’ expression), and tissue space where cells’ colors indicate their task allocation among the archetypes.

**Figure S2:**
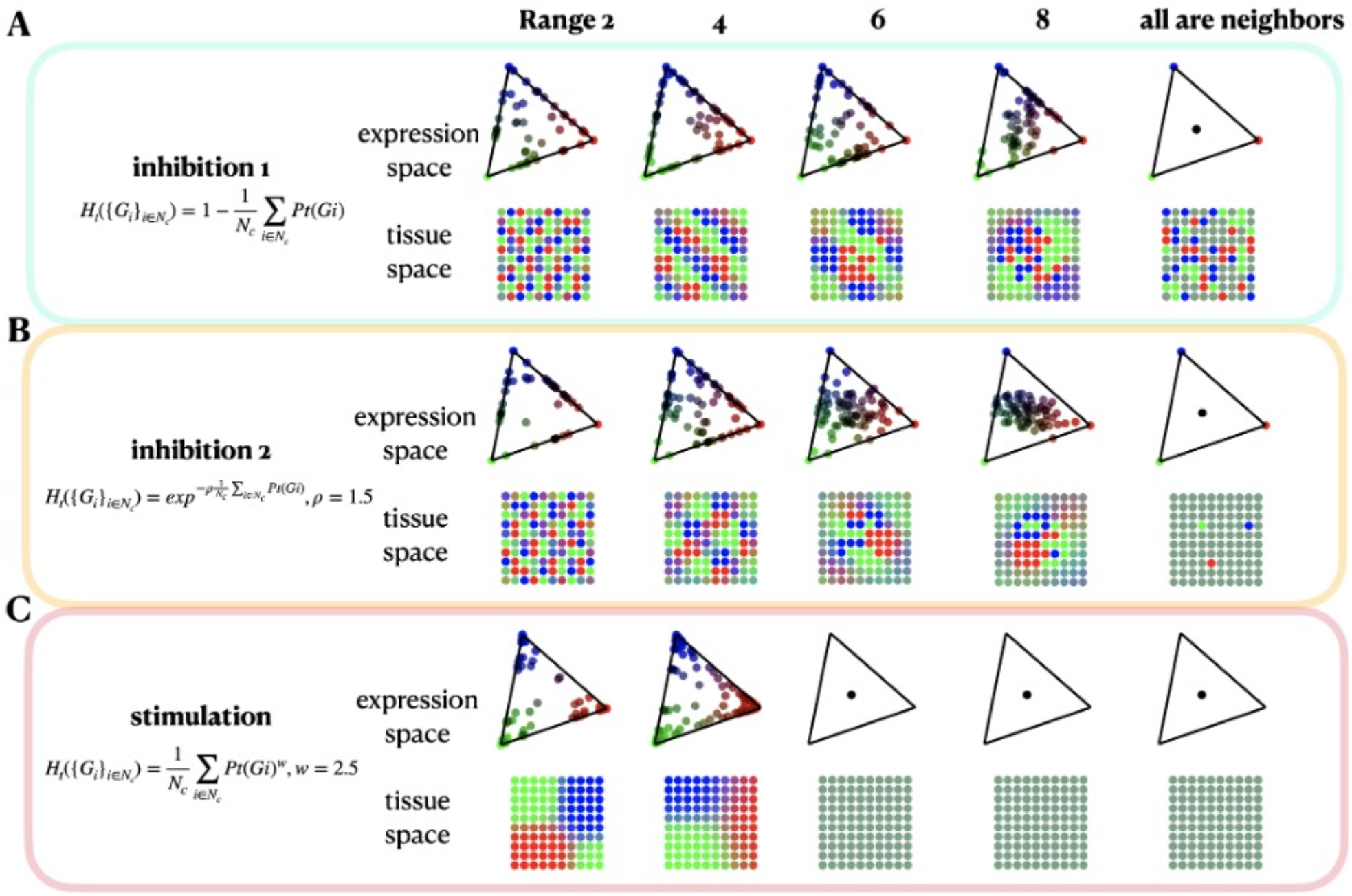
Simulation results for alternative interaction models as a function of the range of interactions. **(A)** The interaction model used throughout the main text. **(B)** Weight of inhibiting interaction decays exponentially with the neighbors’ mean performance in task *t* with rate factor *ρ* = 1.5. **(C)** Stimulating interactions weighted by a mean of the neighbors’ performance to the power of *w* = 2.5.

**Figure S3:**
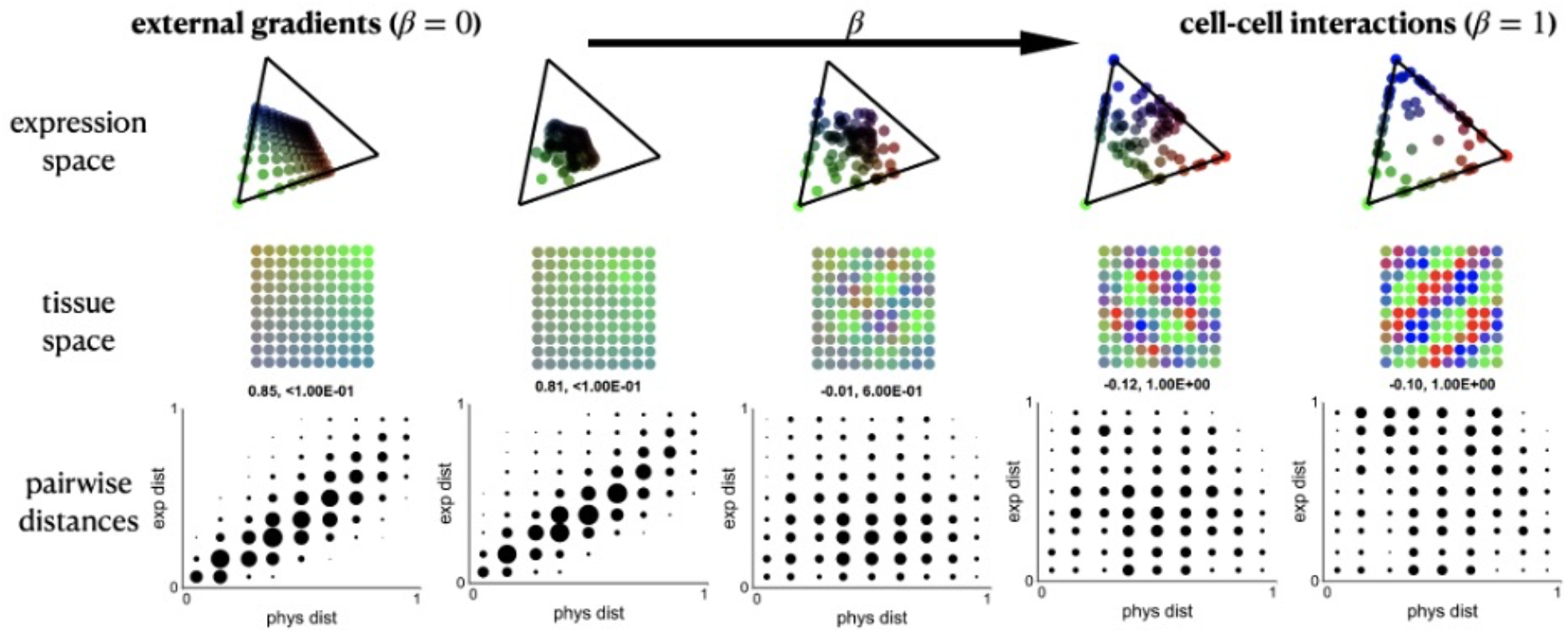
Interpolation between gradients and interactions. *β* = (0,0.5, 0.6, 0.75,1) correspond to columns left to right, for 2D tissue with 3 tasks considering neighborhood of range=4. For each, expression and tissue spaces are plotted. Pairwise distance plots are titled with the Pearson correlation between expression and physical distances, and its p-val.

**Figure S4:**
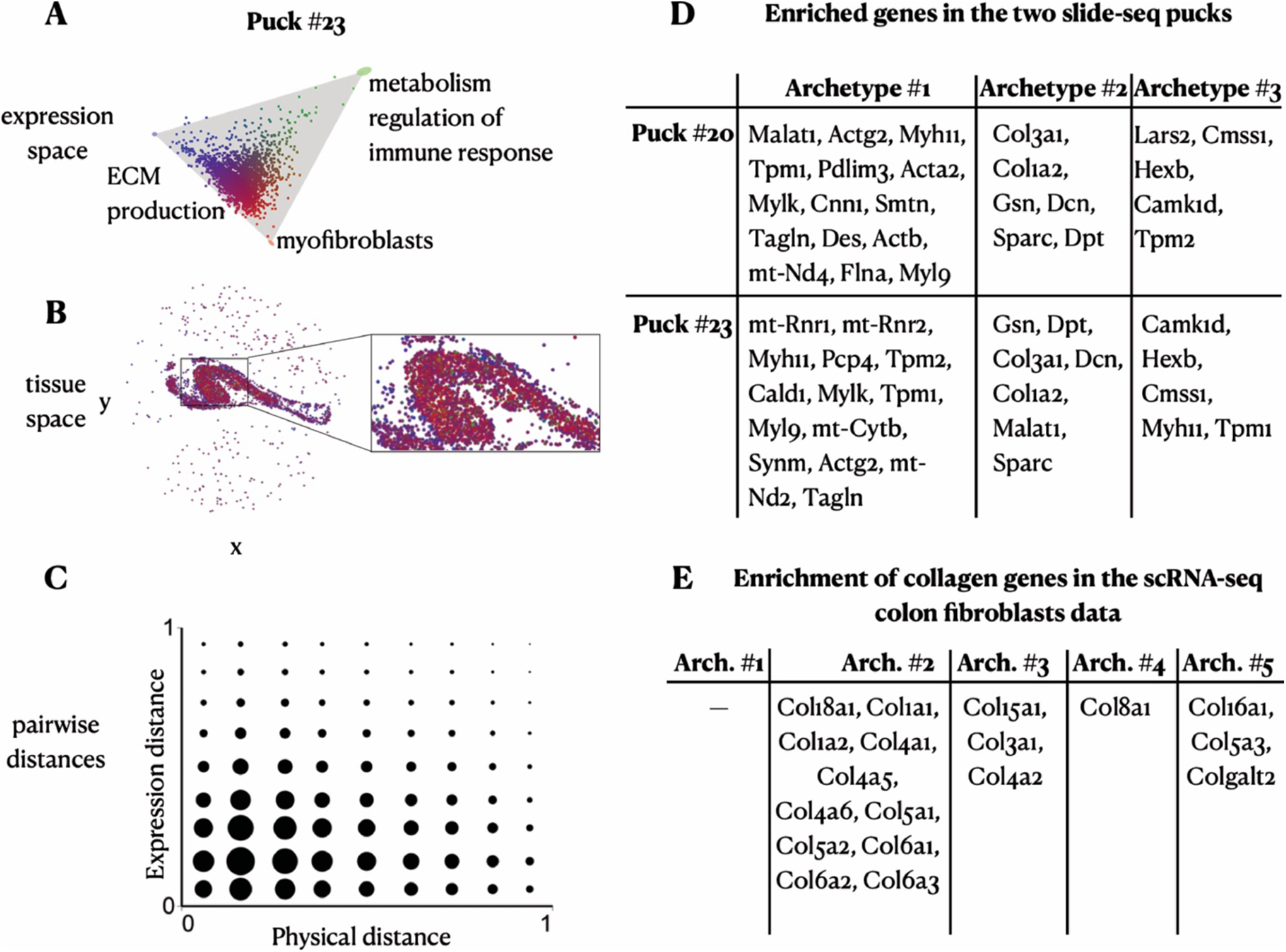
Analysis of additional puck from the Slide-seq data. **(A)** Analysis of puck #23 from the Slide-seq data of colon fibroblasts where cells’ expression profile is projected on the first two PCs, and the cells are colored by their task specializations. **(B)** Spatial coordinates of the cells in tissue space. **(C)** The pairwise distances in physical versus expression space of enterocytes show no correlation (corr=0.004, p-val=0.338). **(D)** A table of enriched genes in the two Slide-seq pucks. **(E)** A table of collagen genes and their enrichment in the scRNA-seq data of mouse colon fibroblasts.

**Figure S5:**
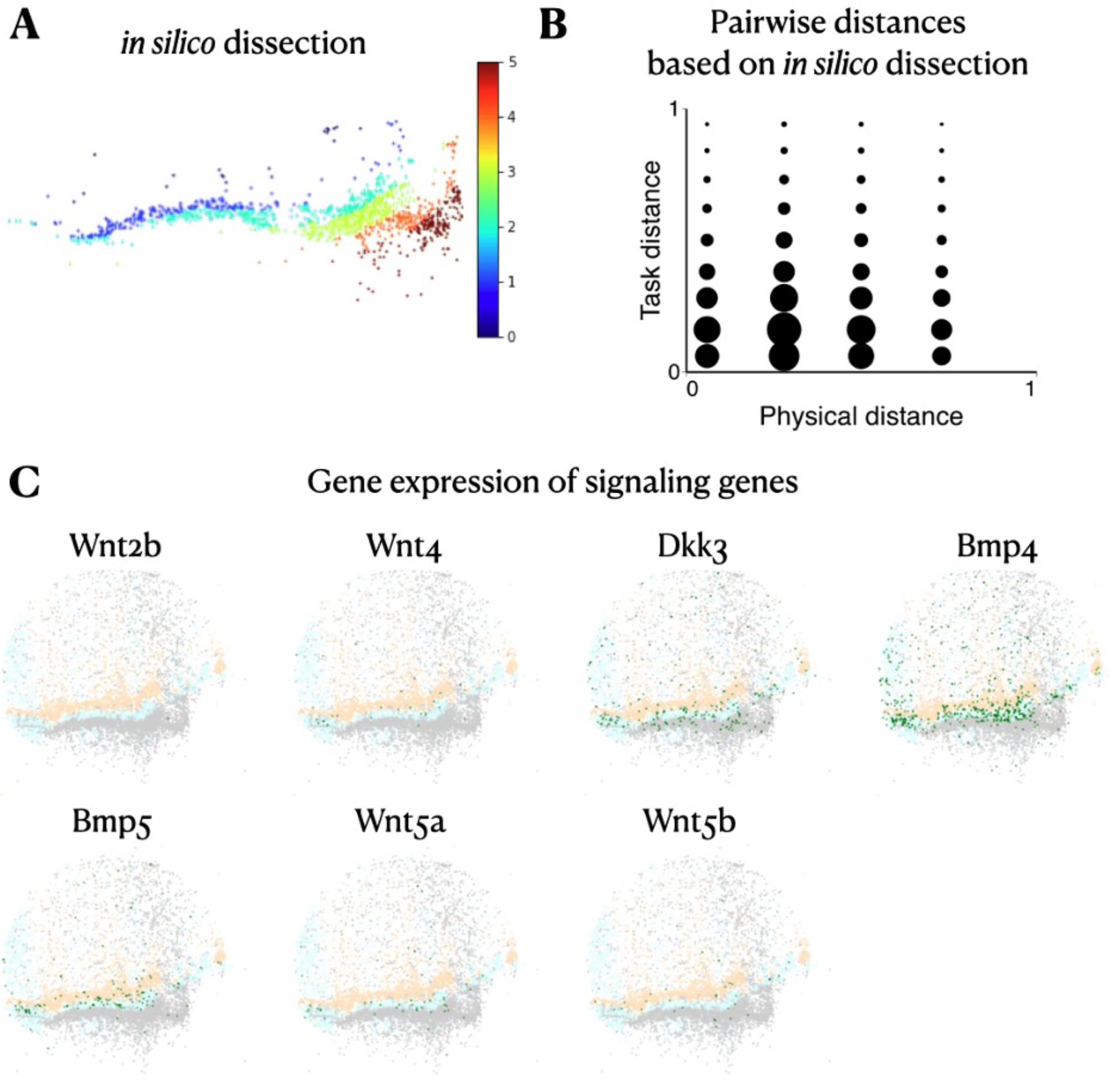
Analysis of in silico dissection and morphogen spatial patterns. **(A)** *In silico* dissection into 6 bins of fibroblasts according to their inferred distance from the deep crypt layer (mimicking how the enterocytes in (Moor et al. 2018) were assayed). **(B)** Considering for each bead its *in silico* bin as its physical location, we compute the corresponding Pearson correlation of pairwise task distances versus physical distances (corr=0.02, p-val=0.037). **(C)** Distribution of signaling genes’ expression across apical-basal colon layers – (light orange) apical plasma membrane, (light blue) deep crypt, (light gray) muscularis. Expression of each gene (in green) is log1p-transformed and truncated at the value of 0.5.

**Figure S6:**
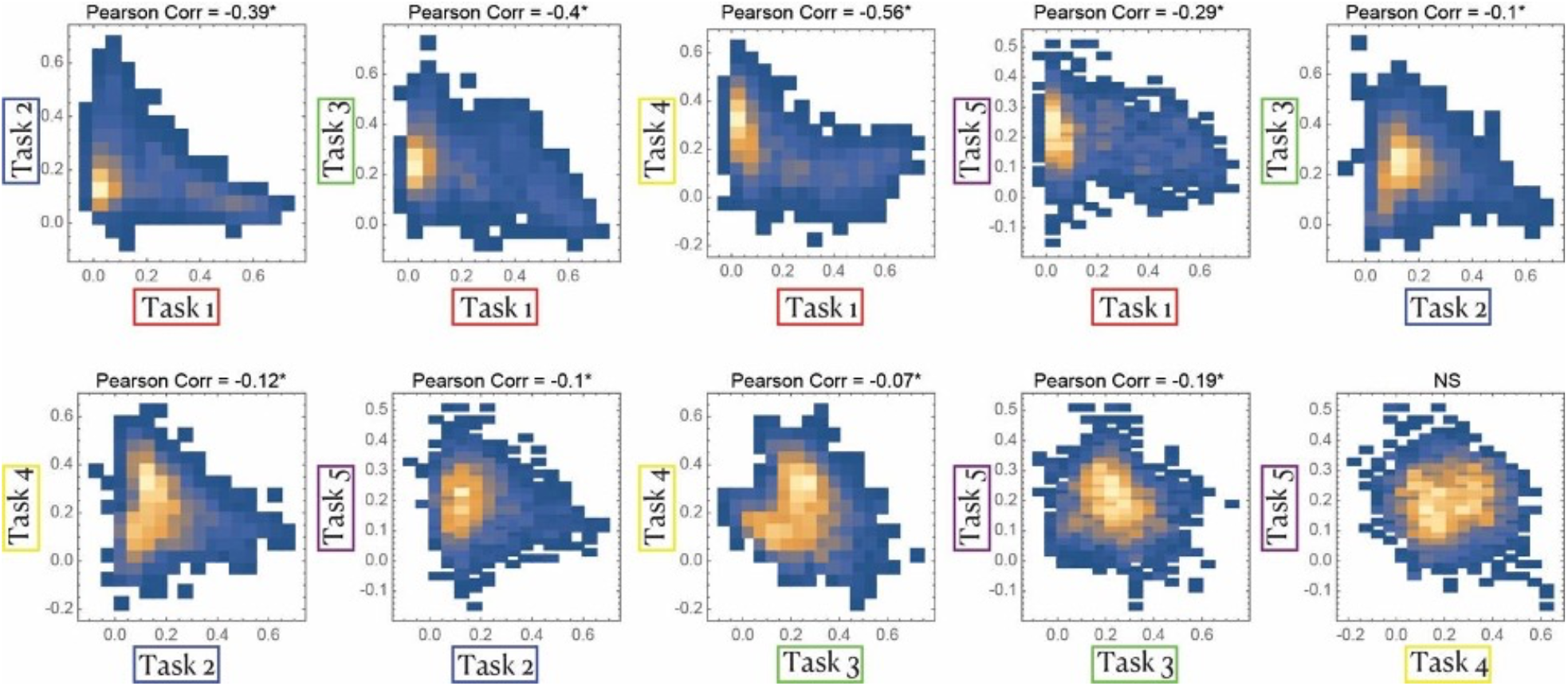
Quantitative analysis of task tradeoffs. Task specialization scores of the Silde-seq fibroblast beads (inferred using the scRNA-seq data analyzed in Figure 4B) are plotted for each pair of archetypes. *p-val<0.001

**Figure S7:**
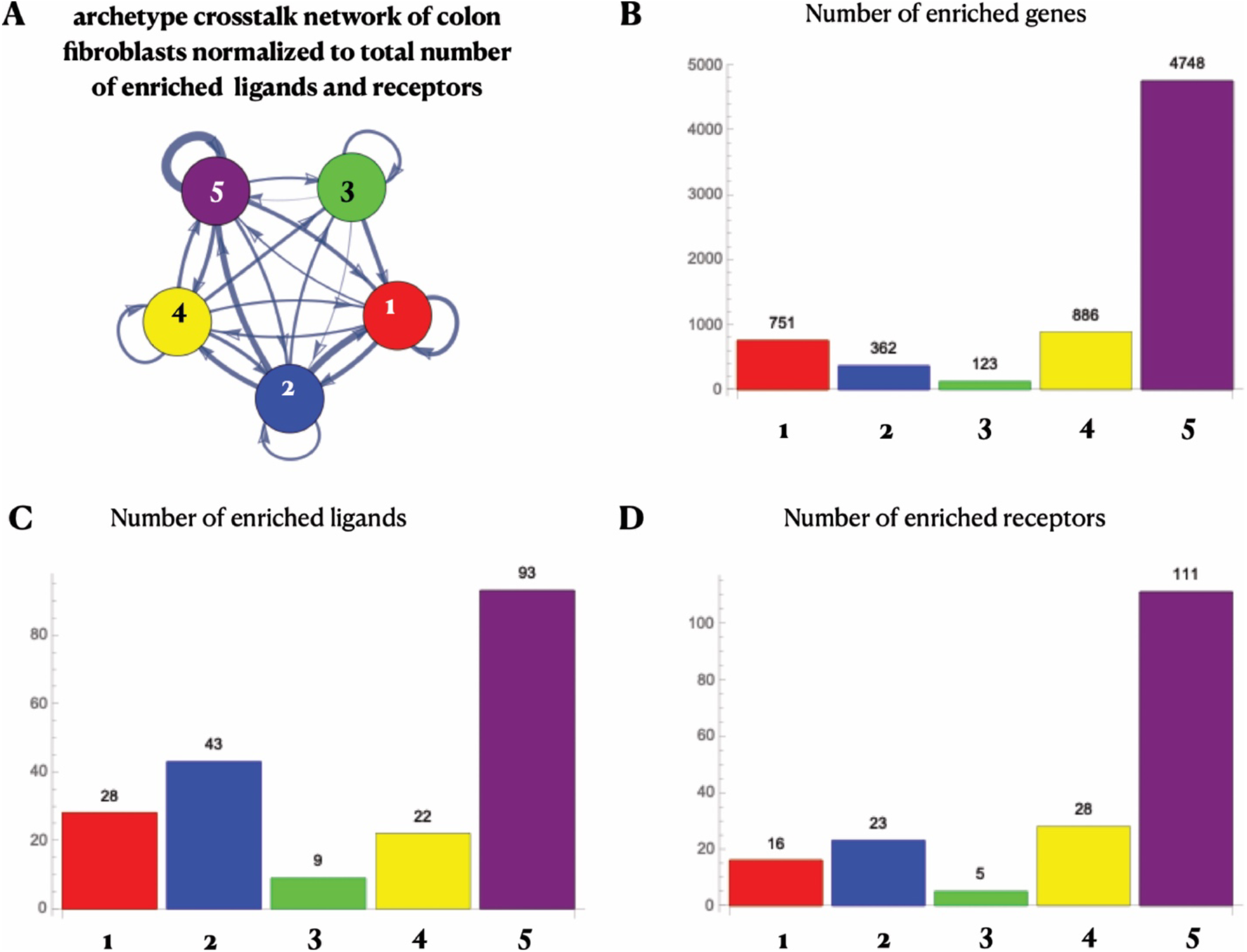
Ligand and receptor enrichments. **(A)** Archetype crosstalk network of the colon fibroblast scRNA-seq data analyzed in **Figure 4B** where the edge weight from archetype A to archetype B is normalized to the sum of the total number of enriched ligands near archetype A and the total number of enriched receptors near archetype B. **(B-D)** Total number of enriched genes (B), ligands (C) and receptors (D) near each of the five colon fibroblast archetypes.

**Figure S8:**
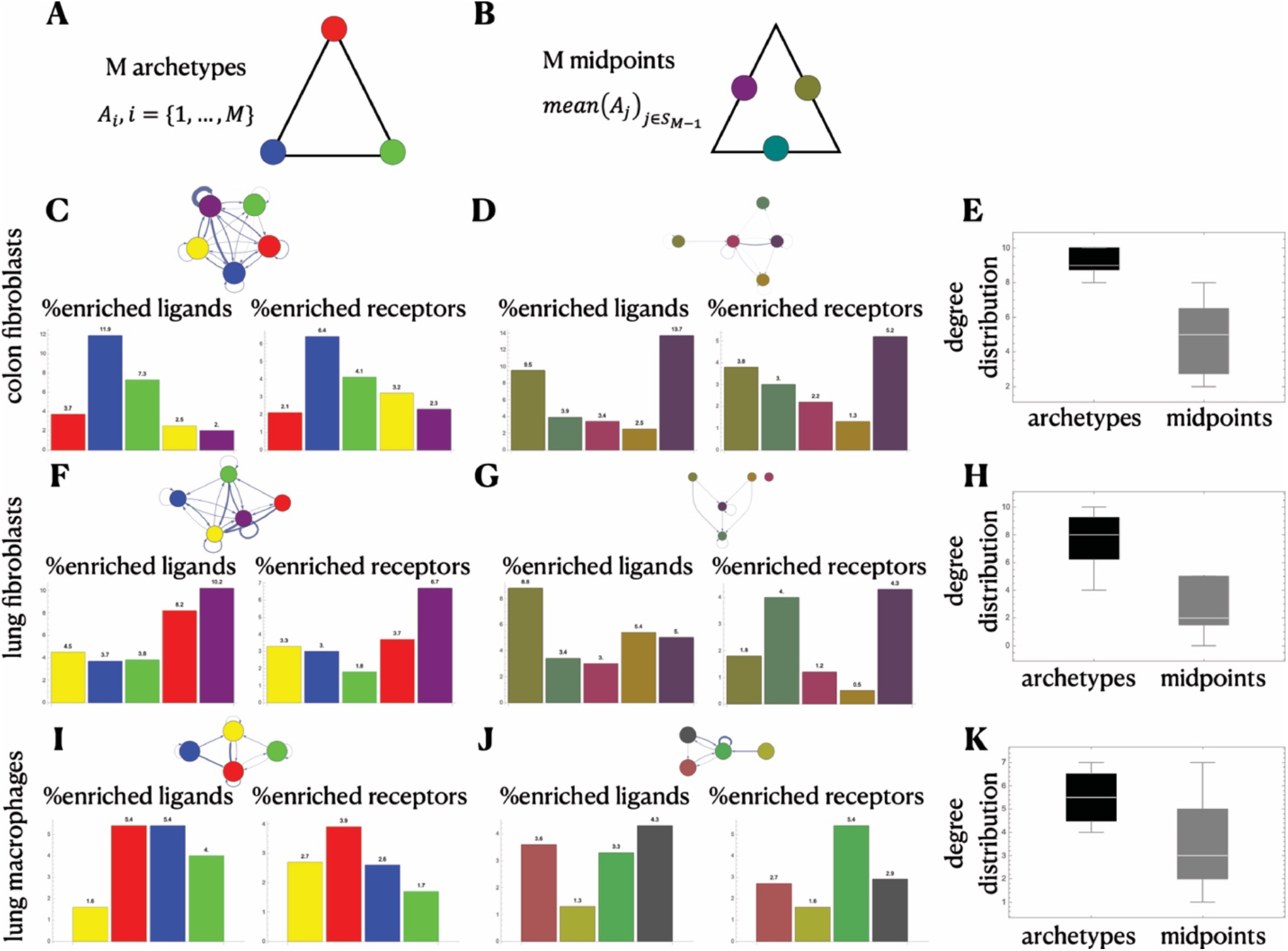
Crosstalk network for archetype midpoints. Comparison of crosstalk networks between the M archetypes **(A)** and between M midpoints **(B). (C-D)** Percentage of enriched ligands and receptors relative to total number of enriched genes near the colon fibroblast archetypes **(C)** and midpoints **(D). (E)** Degree distribution of colon fibroblast archetype crosstalk network (black) and midpoint crosstalk network (gray). The same only for the lung fibroblast data **(F, G, H)** and the lung macrophage data **(I, J, K)**.

## Supplementary information

### Continuum in gene expression space with alternative models of interactions

We discuss in the main text a specific model of inhibiting interactions, where high performance of a task de-prioritizes the task’s allocation in the local region, however, rendered models of interactions can also yield diverse patterns in expression and tissue space and a continuum in expression space. Two examples are demonstrated in Figure S2, changing the inhibiting interaction weight function to decay exponentially with neighbors’ contribution (Figure S2B), or changing the type of interaction to stimulating, that is, neighbors’ performance in task t promote the cell’s performance in the same task (Figure S2C).

### Correspondence between expression and physical distances increases as instructive gradients dominate over interactions (and vice versa)

So far, our model accounted for either external gradients or interactions within the Pareto optimality framework. In practice, however, it is most likely that both mechanisms act simultaneously and jointly shape the gene expression space. We combine the two models and introduce a parameter to interpolate between the effect of external gradients and interactions Figure S3.

### Differences in assays used for colon fibroblasts and intestinal enterocytes do not explain differences in correlation between physical and task pairwise distances

To test whether the difference in spatial resolution between the experimental methods used to assay the colon fibroblasts (assayed with Slide-seq) and the enterocytes along the intestine villus (laser cut into six sections) are the reason for the different resulting correlation values of task and physical distances, we mimic the coarse dissectioning of enterocytes in the analysis of the fibroblasts. To this end, we identify the layers of the colon and infer the distance from the deep crypt layer using TACCO (Mages et al., 2022), and then perform in silico dissection of the fibroblasts according to these distances into six bins (Figure S5A). When using the coarse spatial sectioning of the fibroblasts, we observe a very low Pearson correlation between the task and (new) physical distances (corr=0.02, p-val=0.037, Figure S5B).

### Enrichment of ligand-receptor interactions between archetypes

To test the degree to which the archetypes in the single-cell expression data we analyzed interact with each other through ligand-receptor interactions, we compare the connectivity between the archetypes to the connectivity between a different choice of points in the polytope. To that end, we consider points that are equally distant from each pair of archetypes which we refer to as midpoints. Considering data that fits well within a polytope with M archetypes (Figure S8A), the midpoints are the collection of arithmetic means over every set of M-1 points out the M archetypes (Figure S8B). We used the ParTI package in Matlab to calculate the enriched genes with respect to the midpoints instead of the archetypes. We then constructed the crosstalk network as described above based on ligand-receptor pairs that are enriched near the midpoints. The resulted midpoint crosstalk networks for all three scRNA-seq datasets (Figure S8D, G, J) show that in comparison to the archetype crosstalk networks (Figure S8C, F, I) the number and intensity of links between the vertices declines and not all points interact with each other. The overall number of enriched genes near the midpoints decreases compared to the archetypes. However, the decrease in the crosstalk network connectivity is not due to the decrease in enriched genes. Although the percentage of enriched ligands and receptors out of total number of enriched genes is on the same order of magnitude (Figure S8C-D, F-G, I-J), the number of enriched ligand-receptor *pairs* is significantly lower near the midpoints compared to the archetypes. This is also quantitatively shown when we compute the degree distribution of the two crosstalk networks (Figure S8E, H, K).

